# Diffusional conductance to CO_2_ is the key limitation to photosynthesis in salt-stressed leaves of rice (*Oryza sativa*)

**DOI:** 10.1101/204578

**Authors:** Xiaoxiao Wang, Wencheng Wang, Jianliang Huang, Shaobing Peng, Dongliang Xiong

## Abstract

Salinity significantly limits leaf photosynthesis but the photosynthetic limiting factors in salt- stressed leaves remain unclear. In the present work, photosynthetic and biochemical traits were investigated in four rice genotypes under two NaCl (0 and 150 m*M*) concentration to assess the stomatal, mesophyll and biochemical contributions to reduced photosynthetic rate (*A*) in salt stressed leaves. Our results indicated that salinity led to a decrease in *A*, leaf osmotic potential, electron transport rate and CO_2_ concentrations in the chloroplasts (*C*_c_) of rice leaves. Decreased *A* in salt-stressed leaves was mainly attributable to low *C*_c_, which was determined by stomatal and mesophyll conductance. The increased stomatal limitation was mainly related to the low leaf osmotic potential caused by soil salinity. However, the increased mesophyll limitation in salt stressed leaves was related to both osmotic stress and ion stress. These findings highlight the importance of considering mesophyll conductance when developing salinity-tolerant rice cultivars.

**Abbreviations:**

*A*: photosynthetic rate
*C*_c_, CO_2_: concentration at carboxylation sites
CE: apparent Rubisco activity
Chl: total chlorophyll content
*C*_i_: intercellular CO_2_ concentration
ETR: electron transport rate
F_0_: initial fluorescence of photosystem II in darkness
F_m_: maximum fluorescence of photosystem II
F_v_: maximum variable fluorescence of photosystem II
*F*_v_/*F*_m_: maximum quantum efficiency of photosystem II
*g*_m_: mesophyll conduction
*g*_s_: stomatal conduction
*J*_max_: maximum electron transport rate
K: leaf K content
LMA: leaf mass per area
N: leaf N content
P: leaf P content
OP: osmotic potential
Protein: leaf total soluble protein content
qN: non-chemical quenching efficiency
*R*_d_: day respiration
*R*_dark_: dark respiration
Rubisco: Rubisco content
*V*_cmax_: maximum carboxylation rate
α: leaf light absorptance efficiency
β: the distribution of electrons between PSI and PSII
Γ^*^: CO_2_ compensation point in the absence of respiration
Φ_PSII_: quantum efficiency of photosystem II.

## Introduction

Soil salinity is a global problem that limits crops production worldwide. Rice (*Oryza sativa* L.) is one of the most important cereal crops; however, it has been reported to be very sensitive to salt stress, and it was listed as the most salt-sensitive cereal crop (Munns et al. 2016, Negrão et al. 2011). Salinity reduces rice yield, partially by restraining biomass accumulation which is associated with a decreasing rate of photosynthesis (Moradi and Ismail 2007, Wankhade et al. 2013). A considerable effort has been made, in recent decades, to understand the negative effect of salinity on photosynthesis, but has not yet been fully understood yet. According to the Farquhar model (Farquhar et al. 1980), leaf photosynthesis in C_3_ plants is limited by the capacity of Rubisco to consume RuBP (Rubisco-limited photosynthesis), by the capacity of electron transport and Calvin cycle enzymes to regenerate RuBP (RuBP regeneration-limited photosynthesis) or by the capacity of starch and sucrose synthesis to consume triose phosphates and to regenerate inorganic phosphate for photophosphorylation (Pi regeneration-limited photosynthesis). In general, the Rubisco capacity to consume RuBP is the predominant limitation on photosynthesis at low chloroplasts CO_2_ concentration (*C*_c_) and Rubisco activity; RuBP regeneration limiting related to photosystem electron transport rate (ETR) and the activity of relative enzymes; and Pi regeneration limits photosynthesis under very high *C*_c_. The *C*_c_ is mainly determined by stomatal conductance (*g*_s_), mesophyll conductance (*g*_m_) as well as the Rubisco carboxylation capacity (i.e., *V*_cmax_). Due to the complex responses of leaves to salinity, there is debate over whether the decreased photosynthetic rate (*A*) in salt stressed leaves is primarily limited by *g*_*s*_, *g*_*m*_, electron transport rate (ETR), the activity of relative enzymes (i.e., Rubisco) or a combination of several of these factors (Flexas et al. 2012, Flexas et al. 2016).

A large number of previous studies have described the stomatal limitations in salt-stressed leaves (Centritto et al. 2003, Chaves et al. 2011, Chen et al. 2015, Delfine et al. 1999, Delfine et al. 1998, Moradi and Ismail 2007), because the low leaf water potential introduced by low osmotic potential (termed the ‘osmatic effect’) could cause stomatal closure, while Khan et al. (2015) reported that the decreasing *A* in salt stressed chickpea leaves was predominately caused by photosystem II damage rather than by *g*_s_. However, although the *g*_m_ has rarely been investigated in previous studies, there is no consistent conclusion about mesophyll limitations in salt stressed leaves. Several previous studies observed that both *g*_s_ and *g*_m_ are the primary factors limiting *A* (Centritto et al. 2003, Delfine et al. 1998, Sade et al. 2014). In contrast, other studies have shown that the limitation of *g*_m_ on *A* in salt stressed leaves can be ignored in *Cucumis sativus* (Chen et al. 2015) and *Hordeum vulgare* (Perez-Lopez et al. 2012). These results indicated that the limitations of *g*_m_ on photosynthesis in salt-stressed leaves may be species dependent. Although the *g*_s_ response to salinity in rice has been investigated by many researchers (Moradi and Ismail 2007, Negrão et al. 2011, Wankhade et al. 2013), there have been no previous studies, to our knowledge, investigating the response of *g*m to salinity in rice, which is one of the most salinity sensitive species.

Salinity may also directly inhibit *A* due to the uptake and the accumulation of sodium and chloride in mesophyll tissues (termed as ‘ionic effects’). As the first step of Calvin-Benson cycle and the most abundant protein in C_3_ plants, the content and activity of Rubisco has been suggested as one of the limiting factors involved in reducing *A* under salinity (Delfine et al. 1998, James et al. 2006, James et al. 2002, Yamane et al. 2012). In contrast, Centritto et al. (2003) suggested that the biochemical capacity is not affected by salinity.

The reduction of ETR in salt-stressed leaves which, is often associated with decreases in the actual quantum yield of PSII (Φ_PSII_) and maximal efficiency of PSII photochemistry (F_v_/F_m_) was observed in some species (Koyro et al. 2013, Moradi and Ismail 2007) but not in others (James et al. 2002, Koyro et al. 2013). Similarly, the responses of ETR, Φ_PSII_ and F_v_/F_m_ to salinity were genotype dependent in rice (Moradi and Ismail 2007, Wankhade et al. 2013). The reduction in Φ_PSII_ in some species/genotypes may be due to the salt-induced regulation of energy transduction from the antennae to the reaction centers to prevent photosystem energy surpluses. It was also demonstrated by increased NPQ, an indicator of the excess radiant energy dissipation to heat in the PSII antenna complexes (Murchie and Lawson 2013) under salt-stressed leaves. Moreover, Stepien and Johnson (2009) demonstrated that plastid terminal oxidase acts as an alternative electron sink in *Halophyte thellungiella* a salt tolerant species. Here, we hypothesized that the balance between PSII photochemical activity and the electron requirement for photosynthesis might be broken when CO_2_ concentration in chloroplasts (*C*_c_) decreased due to the reduction of *g*_s_ and *g*_m_ in salinity-sensitive species/genotypes, and this leads to over-excitation and, subsequently, photoinhibition.

In this study, we measured leaf gas exchange and biochemical traits in the model monocot species *Oryza sativa* to reveal the limiting factors of photosynthesis under salinity by using limitation analysis (Buckley and Diaz-Espejo 2014, Grassi and Magnani 2005). The aims of this study as follows: (1) to quantify the limitations of *g*_s_, *g*_m_ and biochemical factors on *A* in salt-stressed leaves; and (2) to test the hypothesis that the decreased *A* in rice is related to photoinhibition under salt stress.

## Materials and Methods

### Plant materials and growth conditions

Rice seeds of four genotypes with different salt tolerances (Xiong *et al.* unpublished data), Liangyoupei 9 (LYP9), N22, Shanyou 63 (SY63) and Texianzhan 25 (TXZ25) were germinated and grown in a nursery for 3 weeks, in a growth chamber (Model GR48, Conviron, Controlled Environments Limited, Winnipeg, MB, Canada). In the chamber, the air temperature was set at 28°C/22°C (day/ night), with a relative humidity at 70% and PPFD at 600 molm^−2^ s^−1^ with a 12 h: 12h light/dark regime. The plants were then transplanted into 11-l plastic pots containing 10 kg of soil with a density of three plants per pot in the same chamber. Before transplanting, 7.0 g of compound fertilizer (N: P_2_O_5_: K_2_O=16: 16: 16%, Batian Ecological Engineering Limited, Shenzhen, China) per pot was mixed into the soil, and, 30 days after transplanting, 1.3 g of urea per pot was top-dressed. For each genotype, 10 pots were grown and random arranged. To avoid water stress, at least a 2-cm water layer was maintained. Seven weeks after transplanting, half of the pots of each genotype were irrigated with 1 l of 150 m*M* NaCl solution every two days for one week. All of the measurements were performed in fully expanded young leaves.

### Gas exchange and chlorophyll fluorescence measurements

Gas exchange was measured in a growth chamber between 8:30 and 16:00 and measurements were carried out on the newly and fully expanded leaves of three plants in each treatment. Gas exchange was measured using a Licor-6400 portable photosynthesis system equipped with a Li-6400-40 chamber (LI-COR Inc., Lincoln, NE). In the leaf chamber, the PPFD was maintained at 1200 jumol m^−2^ s^−1^, a leaf-to-air vapor pressure deficit (*VPD*) at 1.5-2.0 kPa, and a CO_2_ concentration adjusted to 400 jumol m^−2^ s^−1^ with a CO_2_ mixer. The block temperature during measurements was maintained at 28°C. After equilibration to a steady state (usually more than 20 min after clamping the leaf), the gas exchange parameters, steady-state fluorescence (*F*_s_) and maximum fluorescence (*F*_m_‵) were recorded. The actual photochemical efficiency of photosystem II (Φ_PSII_) was calculated as follows:

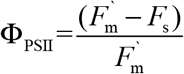

The electron transport rates (ETR) were computed as follows:

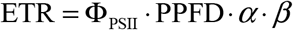

where *α* is the leaf absorbance, and *β* represents the distribution of electrons between PSI and PSII.

Five days after the NaCl treatment, the light response curves were performed under low O_2_ concentration (< 2%) to estimate *α* and *β*. The gas exchange system was switched to low O_2_ concentration (< 2%) by injecting pure N_2_. Simultaneous measurements of light response curves and chlorophyll fluorescence were then performed. During the measurements, the chamber conditions were the same as those described above, except that PPFD was controlled across a gradient of 2000, 1500, 1200, 1000, 800, 600, 400, 200, 100, 0 and 1200 μmol m^−2^ s^−1^. After reaching a steady state, the parameters of gas exchange and chlorophyll fluorescence were simultaneously recorded. The slope of the relationship between Φ_PSII_ and 4ΦCO_2_ (the quantum efficiency of CO_2_ uptake) is considered to be the value of *α·β* (Valentini et al. 1995). There were no differences in *α·β* values between the control and the salt stressed leaves (Fig. S1 B), thus the average value for all the genotypes were used in the current study.

The mesophyll conductance of CO_2_ (*g*_m_) was calculated based on the variable *J* method described in (Harley et al. 1992). In this method, the CO_2_ concentration in the chloroplast (*C*_c_) was calculated as follows:

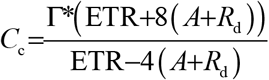

where Γ^*^ represents the CO_2_ compensation point in the absence of respiration and *R*_d_ is the day respiration, which was assumed to be half of the dark respiration rate (*R*_dark_). Γ* is related to the Rubisco specific factor (*S*_C/O_), which is relatively conserved under a given temperature condition. In the present study, rice *S*_C/O_ at 28°C was obtained from Hermida-Carrera et al. (2016). Then, *g*_m_ was calculated as follows:

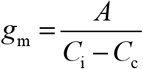

where *C*_i_ represents the intercellular CO_2_ concentration.

After 7 days of salt treatment, the CO_2_ response curves (*A-C*_i_ curves) were measured with two Li-6400 portable photosynthesis system equipped with a Li-6400-40 chamber in three days. The CO_2_R was set at 400, 300, 200, 150, 100, 50, 0, 400, 600, 800, 1000, 1500, 2000 and 400 μmol mol^−1^, the photosynthetic photon flux density (PPFD) was set as 1200 μmol m^−2^ s^−1^ with a 10:90 blue: red light, the flow rate at 150 μmol s^−1^ and the relative humidity at 65(±5)%. When the stomatal conductance was stable (less than 5% variation during 10 min), the automatic program was run. For each step, 4 to 5 min were waited. The dead rice leaves (obtained by heating the leaves until no variable chlorophyll fluorescence was observed) were used to estimate the leakage effects of the chamber under different CO_2_ concentrations (Flexas et al. 2007, Xiong et al. 2015b). In the current study, the sum of the photorespiration and mitochondrial respiration in the light (*R*_L_) was calculated by extrapolating the *A-C*_i_ curve to *C*_i_ = 0 (Escalona et al. 1999, Flexas et al. 2002).

Seven days after NaCl treatment, the *g*_m_ was calculated with two methods: Harley’s method (Harley et al. 1992) and Ethier’s method (Ethier and Livingston 2004). The method of Ethier and Livingston (2004) uses only gas exchange measurements, by adjusting the non-linear model of Farquhar et al. (1980) to extract the *g*_m_. In the present study, some NaCl treatment leaves did not reach satisfactory results by using Ethier’s method.Therefore, we used the values obtained by Harley’s method to compare with other parameters in the manuscript, while a good correlation was obtained between the two estimates of *g*_m_ considering the data averaged per treatments (*R*^2^=0.86; *P*< 0.001; Fig. S1 B).

Dark respiration (*R*_dark_) was measured by Li-Cor 6400 after *A-C*_i_ curves were performed. Before the *R*_dark_ was measured, rice plants were acclimatized to darkness for at least 2.0 h. In the Li-Cor leaf chamber, the ambient CO_2_ concentration was adjusted to 400 μmol mol^−1^ using a CO_2_ mixture, the block temperature was maintained at 28 °C, the PPFD was 0 μmol m^−2^ s^−1^, the leaf-to-air vapor pressure deficit (VPD) was between 1.1 and 1.5 kPa, and the flow rate was 100 mol s^−1^. After the leaf reached a steady state, usually after 10 min, gas exchange parameters were recorded.

### Photosynthetic limitation analysis

Limitation analysis is a helpful tool to quantify the stress effects of changes in various factors on *A* (Buckley and Diaz-Espejo 2014, Grassi and Magnani 2005), and it has been widely used in recent years (Chen et al. 2015, Flexas et al. 2009, Galle et al. 2011, Galle et al. 2009, Tosens et al. 2015). Relative photosynthetic limitations including stomatal (*l*_s_), mesophyll (*l*_s_) and biochemical (*l*_b_) relative limitations were calculated according to Grassi and Magnani (2005).

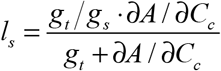

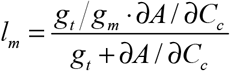

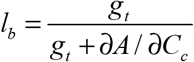

To assess the effects of salinity on changes in photosynthetic limitations in each genotype and treatment duration, the relative limitations were linked to overall changes in *A*:

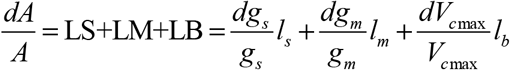

where LS, LM and LB are the reduction fractional limitation in *A* caused by reduction in stomatal conductance, mesophyll conductance and biochemistry, respectively. In the current study, the photosynthetic parameters of control were defined as the references.

### Chlorophyll fluorescence

Chlorophyll fluorescence parameters were measured at pre-dawn to investigate the F_v_/F_m_. A portable pulse amplitude modulation fluorescence instrument (PAM 2000, Walz, Effeltrich, Germany) was used. A measuring light of approximately 0.5 μmol photons m^−2^ s^−1^ was set to a frequency of 600 Hz to determine the background fluorescence signal (F_o_) as well as the maximum fluorescence (F_m_), and the F_v_ was calculated as F_v_=F_m_−F_o_.

### Determination of N, P, Na and K content

After seven-days salt treatment, the fully expanded young leaves were sampled after taking a picture for leaf area estimation, and were dried under 80°C to a constant weight. The dry samples were digested by the mocro-Kjeldahl method (Xiong et al., 2015c). The N and P concentrations were measured with a discrete wet chemistry analyzer (SmartChem 200, AMS-Westco, Rome, Italy). The Na and K concentration were measured by an atom absorption spectrometer (PinAAcle 900T, Perkin Elmer, Waltham, MA). The leaf area was measured by using image J software (National institute of Health, Bethesda, MD).

### Determination of the total soluble protein, Rubisco and Chlorophyll content

Leaf samples were harvested in the morning of the seventh-day after NaCl treatment, and immersed in liquid nitrogen. The samples were stored at −80°C until the solution protein and Rubisco concentration were measured. The frozen leaf sample was ground in liquid nitrogen and homogenized in ice in an extraction buffer containing 50 mM Tris-HCl buffer (pH 8.0), 5 mmol β-mercaptoethanol, and 12.5% glycerol (v/v). After centrifuging, the supernatant fluid was used as a total solution protein as well as for a Rubisco content analysis (Xiong et al. 2015b, Xiong et al. 2015c). The Rubisco samples were loaded onto SDS-PAGE containing a 12.5% (w/v) polyacrylamide gel. After electrophoresis (DYY-11, Beijing Liuyi Instrument Factory), the gels were washed with deionized water several times and then dyed in 0.25% commassie blue staining solution for 9 h and decolorized until the background was colorless. Then, the Rubisco was transferred into a 5-ml cuvette with 1.5 ml of formamide and washed in a 50°C water bath at room temperature for 8 h. The washed solutions were measured at 595 nm (Infinite M200, Tecan U.S., Inc) using the background glue as a blank, and bovine serum albumin (BSA) as the standard protein.

### Osmotic potential measurements

The fully expanded young leaves were sampled in the morning of the seven-day after NaCl treatment. The leaf samples were immersed in liquid nitrogen and then stored at −80°C until measured. The leaf osmotic potential was measured by using a vapor pressure osmometer (VAPRO 5520, Wescor Inc., Logan, Utah).

### Statistical analysis

ANOVA with a post hoc Tukey HSD test was used to test the differences and interactions in the measured traits among genotypes and treatments. Regression analyses were performed with mean values to test the correlations between parameters. Regressions were fitted with linear model, except in Fig. 5, which fitted by a power model (y=ax^b^). Regression lines were shown for *P* < 0.05. All of the analyses were performed in R version 3.3.1 (https://cran.r-project.org).

## Results

We observed substantial variations in the responses of chemical composition, photosynthetic traits, chlorophyll fluorescence as well as the osmotic potential of salinity in rice (Fig. 1). The salinity responses of those traits also varied with genotype; overall, N22 was more tolerant of salinity than the other three genotypes.

**Fig. 1.**
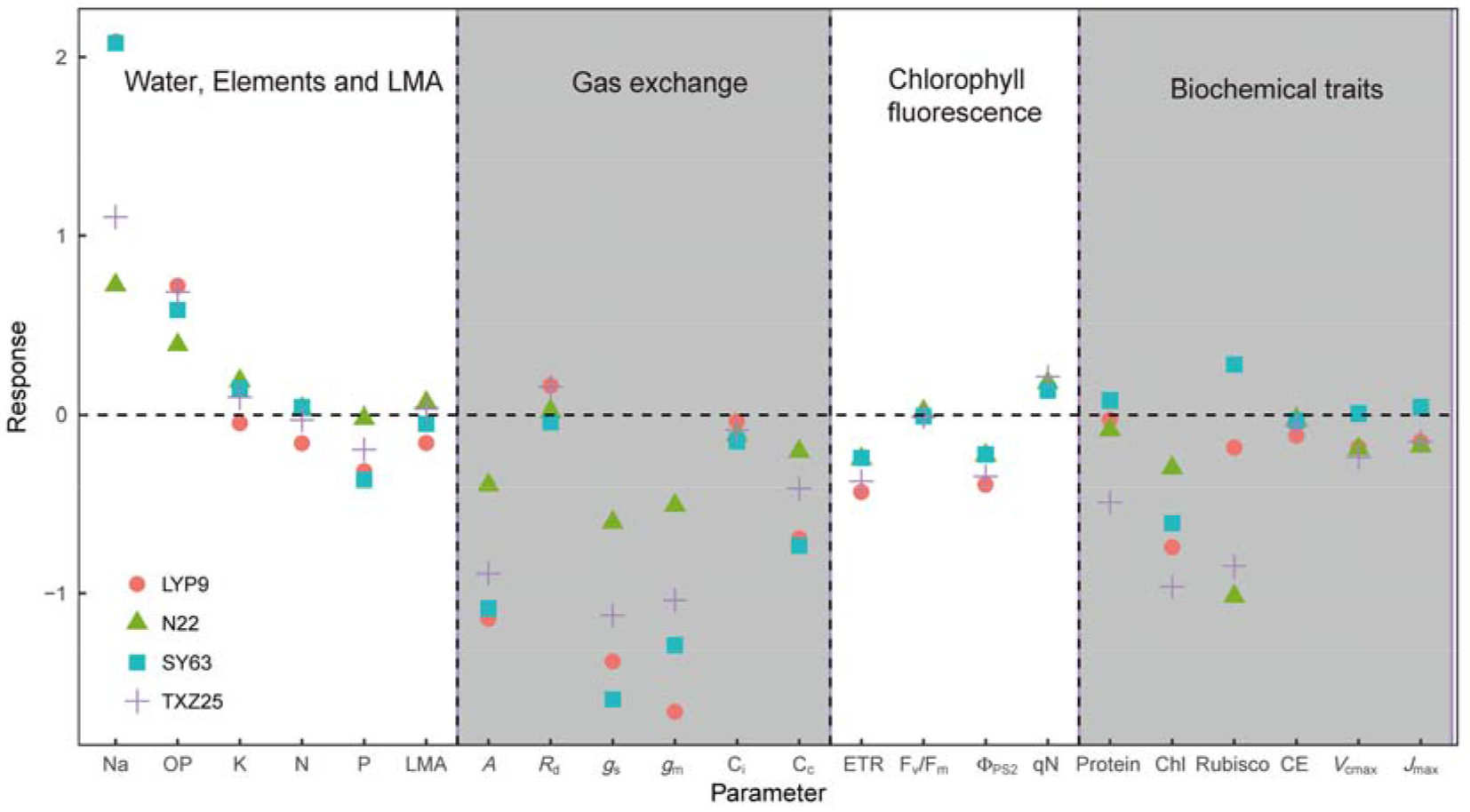
Response of variation to 7-days salinity in four rice genotypes. The responses were calculated by ln(X_T_/X_CK_), where the X_T_ and X_CK_ represent the mean values of the parameter under NaCl treatment and control, respectively.

### Effects of salinity on leaf biochemical parameters

Overall, the leaf Na^+^, P and K content in salt-stressed rice increased significantly after seven days of NaCl treatments, while a substantial genetic variation was observed (Table 1; Fig. 1). There was no difference in the LMA and leaf N content between the control and salinity treatment, despite a significant variation among genotypes. Across all four genotypes, salt stress decreased the total leaf solution protein and Rubisco content, whereas both the total leaf solution protein and Rubisco content increased in SY63.

**Table 1.** Effects of salinity on leaf mass per area and leaf biochemical traits in four rice genotypes. N, nitrogen content; P, phosphorus content; K, potassium content; Na, sodium content; protein, solution protein content; Rubisco, Rubisco content; T, treatment and G, genotype. Different lower case letters indicate a significant difference between treatment and control for a given genotype (*P* < 0.05). ns, no significant; \**P* < 0.05; \*\**P* < 0.01; and \*\*\**P* < 0.001.

| Genotype | Treatment | LMA (g m<sup>-2</sup>) | N (g m<sup>-2</sup>) | P (mg m<sup>-2</sup>) | K (g m<sup>-2</sup>) | Na (mg m<sup>-2</sup>) | Protein (g m<sup>-2</sup>) | Rubisco (g m<sup>-2</sup>) |
| --- | --- | --- | --- | --- | --- | --- | --- | --- |
| LYP9 | CK | 60.6±4.0 | 1.41±0.12 | 99.0±4.4 | 1.14±0.09 | 5.7±1.2 | 4.41±0.05 | 1.82±0.64 |
|  | NaCl | 51.8±6.4 | 1.20±0.11 | 72.1±8.3 | 1.08±0.18 | 46.1±17.0 | 4.30±0.28 | 1.51±0.46 |
| N22 | CK | 37.7±1.2 | 1.09±0.06 | 66.7±4.0 | 0.57±0.02 | 3.7±1.3 | 3.41±0.21 | 1.65±0.26 |
|  | NaCl | 40.2±2.3 | 1.12±0.03 | 65.2±1.5 | 0.69±0.03 | 7.6±2.2 | 3.13±0.20 | 0.6±0.10 |
| SY63 | CK | 53.6±2.9 | 1.44±0.11 | 112.9±9.1 | 0.72±0.07 | 5.5±1.4 | 3.48±0.44 | 1.13±0.32 |
|  | NaCl | 50.7±4.6 | 1.49±0.15 | 78.3±8.0 | 0.83±0.08 | 43.8±25.1 | 3.76±0.18 | 1.49±0.40 |
| TXZ25 | CK | 40.1±3.7 | 1.29±0.12 | 90.3±9.7 | 0.55±0.07 | 6.5±1.1 | 4.37±0.42 | 1.55±0.47 |
|  | NaCl | 41.6±3.3 | 1.25±0.09 | 74.2±8.5 | 0.61±0.06 | 19.6±4.0 | 2.67±0.20 | 0.66±0.21 |
| Significance | T | ns | ns | * | * | ** | * | *** |
|  | G | * | * | * | * | * | ns | ns |
|  | T×G | ns | ns | * | * | ** | * | *** |

### Gas exchange and osmotic potential

*A*, *g*_s_, *g*_m_ and ETR in salt-stressed rice leaves declined rapidly after starting the NaCl treatment (Fig. S2). *A* decreased by approximately 30% in the salt-stressed leaves of SY63 and TXZ25, but no response in N22 on the first day after NaCl treatment. After three days of NaCl treatment, the *A* in salt stressed leaves of all the four genotypes decreased. A Similar response pattern was found in *g*_s_, *g*_m_ and ETR (Fig. S2). After seven days of NaCl treatment, the biggest decline of *A* was in the salt-stressed leaves of LYP9 (72%) and the mildest decline occurred in N22 (38%) (Fig. S2). The gas exchange and osmotic potential parameters after seven days of NaCl treatment are shown in Table 2. Overall, substantial variation in *A* of rice leaves was found among genotypes as well as in the NaCl treatments. Similar to *A*, both *g*_s_ and *g*_m_ decreased significantly in the salt stressed leaves of LYP9, SY63 and TXZ25, but not in N22 (Fig. 2). Across the genotypes and NaCl treatments, *A* was tightly correlated with *g*_s_ (*R*^2^=0.91; *P* < 0.001) and g_m_ (*R*^2^=0.98; *P* < 0.001). There was no significant difference of *R*_dark_ and CE among genotypes and NaCl treatments. While no genetic variation in *C*_i_, *C*_c_, *V*_cmax_, *J*_max_ and osmotic potential were found, salinity significantly decreased those parameters.

**Table 2.** Effects of salinity on leaf physiological traits in four rice genotypes. A, light-saturated photosynthetic rate; *R*_dark_, dark respiration; *C*_i_ intercellular CO_2_ concentration; *C*_c_, CO_2_ concentration at chloroplasts, *V*_cmax_, maximum carboxylation rate; *J*_max_, maximum electron transport rate; and OP, osmotic potential. Different lower case letters indicate a significant difference between treatment and control for a given genotype (*P* < 0.05). ns, no significant; \**P*<0.05; \*\**P*<0.01 and \*\*\**P*<0.001.

| Genotype | Treatment | $A$ ($\mu\text{mol m}^{-2} \text{s}^{-1}$) | $R_{\text{dark}}$ ($\mu\text{mol m}^{-2} \text{s}^{-1}$) | $C_i$ ($\mu\text{mol mol}^{-1}$) | $C_c$ ($\mu\text{mol mol}^{-1}$) | $V_{\text{cmax}}$ ($\mu\text{mol m}^{-2} \text{s}^{-1}$) | $J_{\text{max}}$ ($\mu\text{mol m}^{-2} \text{s}^{-1}$) | CE ($\text{mol m}^{-2} \text{s}^{-1}$) | OP (MPa) |
| --- | --- | --- | --- | --- | --- | --- | --- | --- | --- |
| LYP9 | CK | 11.58±2.98 | 0.672±0.010 | 302±26 | 189±21 | 114.2±31.2 | 123.7±21.3 | 0.11±0.05 | -0.98±0.12 |
|  | NaCl | 3.34±0.67 | 0.790±0.048 | 291±20 | 93±10 | 95.1±3.0 | 106.4±14.3 | 0.12±0.01 | -2.02±0.08 |
| N22 | CK | 8.53±2.25 | 0.500±0.017 | 283±23 | 131±32 | 104.5±2.8 | 110.5±6.9 | 0.14±0.03 | -1.30±0.18 |
|  | NaCl | 5.58±1.54 | 0.510±0.016 | 253±31 | 111±12 | 86.3±8.2 | 92.7±14.0 | 0.13±0.01 | -1.92±0.20 |
| SY63 | CK | 11.18±0.87 | 0.778±0.027 | 291±9 | 189±39 | 132.1±2.4 | 135.7±20.4 | 0.15±0.02 | -1.15±0.05 |
|  | NaCl | 3.42±1.44 | 0.747±0.015 | 251±34 | 84±6 | 132.9±22.9 | 141.4±8.0 | 0.15±0.04 | -2.06±0.19 |
| TXZ25 | CK | 9.65±0.95 | 0.574±0.030 | 265±8 | 122±24 | 144.7±1.4 | 148.3±20.5 | 0.16±0.02 | -0.94±0.08 |
|  | NaCl | 3.65±0.19 | 0.669±0.041 | 243±28 | 83±4 | 114.0±6.5 | 127.4±17.5 | 0.15±0.01 | -1.88±0.09 |
| Significance | T | *** | ns | * | ** | * | * | ns | *** |
|  | G | * | ns | ns | ns | ns | ns | ns | ns |
|  | T×G | ** | ns | ns | ** | * | * | ns | ns |

**Fig. 2.**
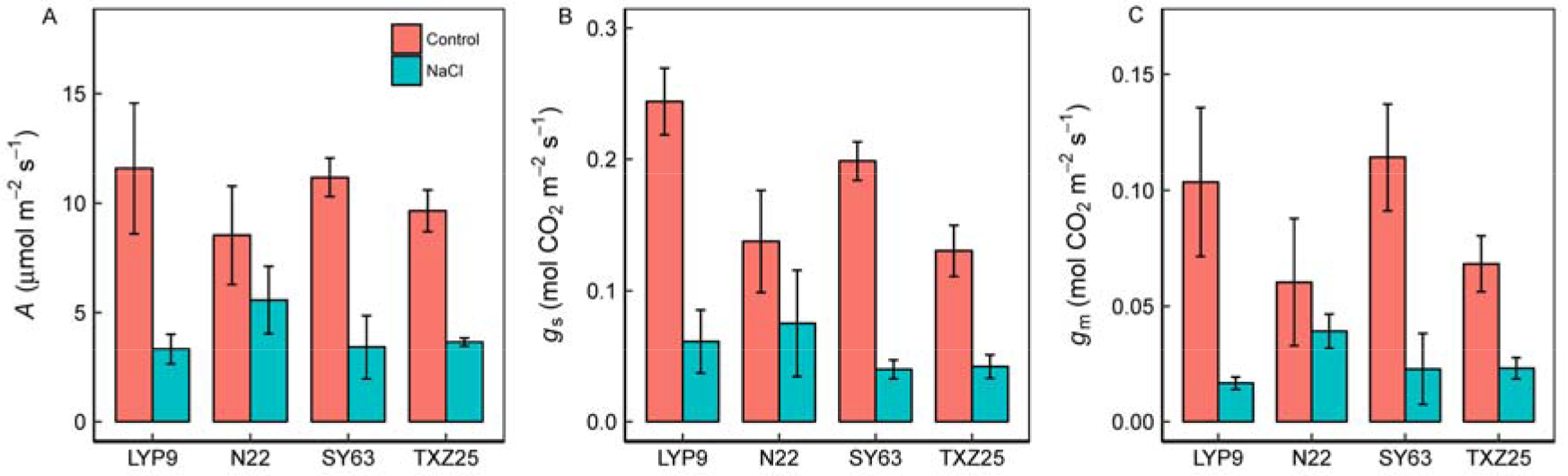
Effects of 7-days salt treat on (A) Light saturated photosynthetic rate (*A*), (B) stomatal conductance (*g*_s_) and (C) mesophyll conductance (*g*_m_) of four rice genotypes. Bars represent the mean ± se of at least three replicates. ns, no significant; *, *P*<0.05; **, *P*<0.01; and ***, *P*<0.001.

**Fig. 3.**
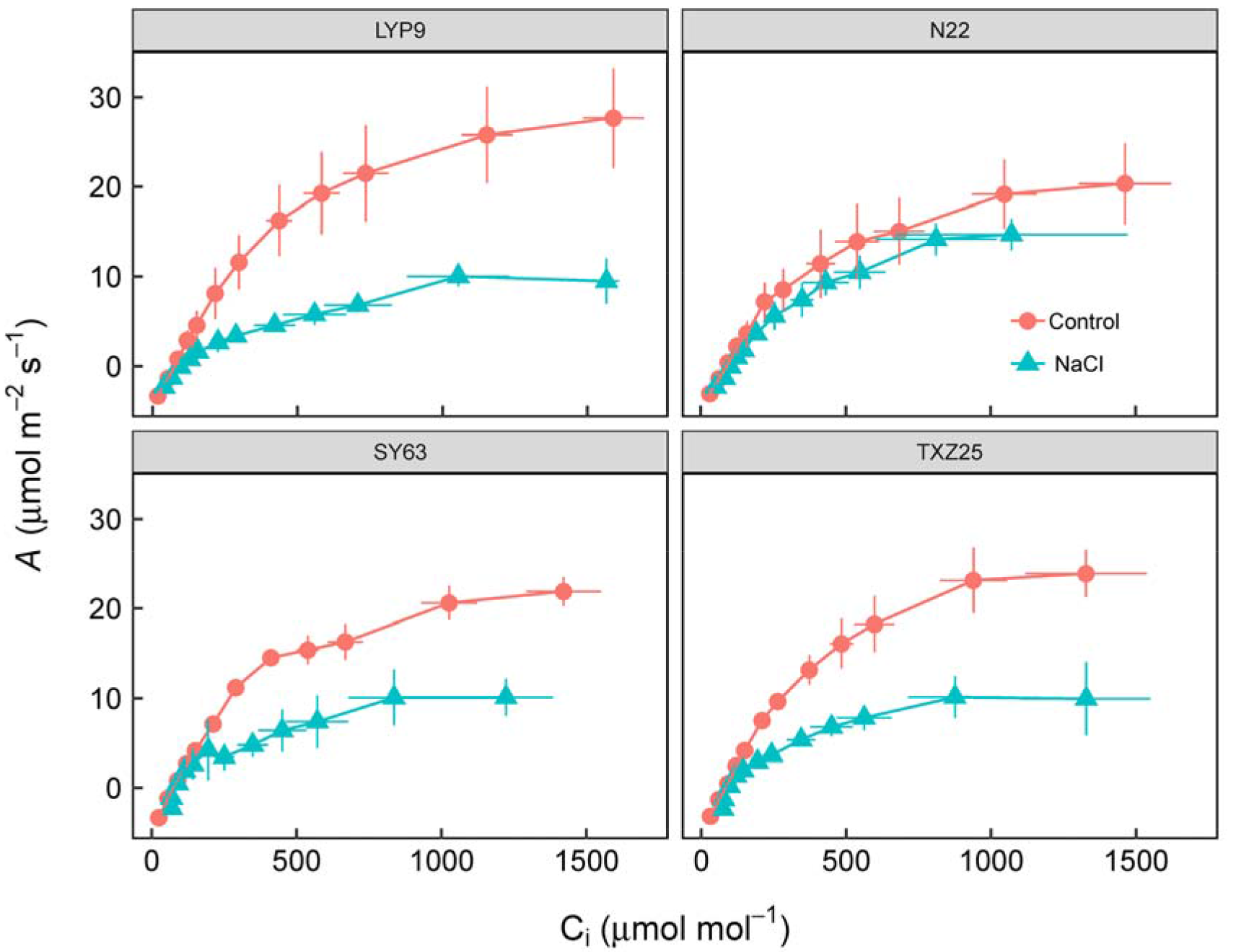
Responses of light-saturated photosynthetic rate (*A*) to intercellular CO_2_ concentration (C_i_) in four rice genotypes. Points represent the mean ± se of at least three replicates.

Both ETR and ΦPSII trended lower in salt-stressed leaves than in the control (Fig. 4), although only LYP9 and TXZ25 showed statistical significance (Fig. 4). However, F_v_/F_m_ showed no difference between the control and salinity in all of the estimated genotypes (Fig. 4B). in contrast, qN exhibited an increasing tendency in salt stressed leaves, while only TXZ25 exhibited a significant increase. The ratio of ETR/A varied widely with varying C_c_ across four genotypes and two NaCl treatments (Fig. 5); however, the ratio of ETR/(*A*+*R*_L_) exhibited constant with varying *C*_c_. When, *C*_c_ was lower than 100 μmol mol^−1^, the ratio of ETR/*A* increased fast with decreasing *C*_c_.

**Fig. 4.**
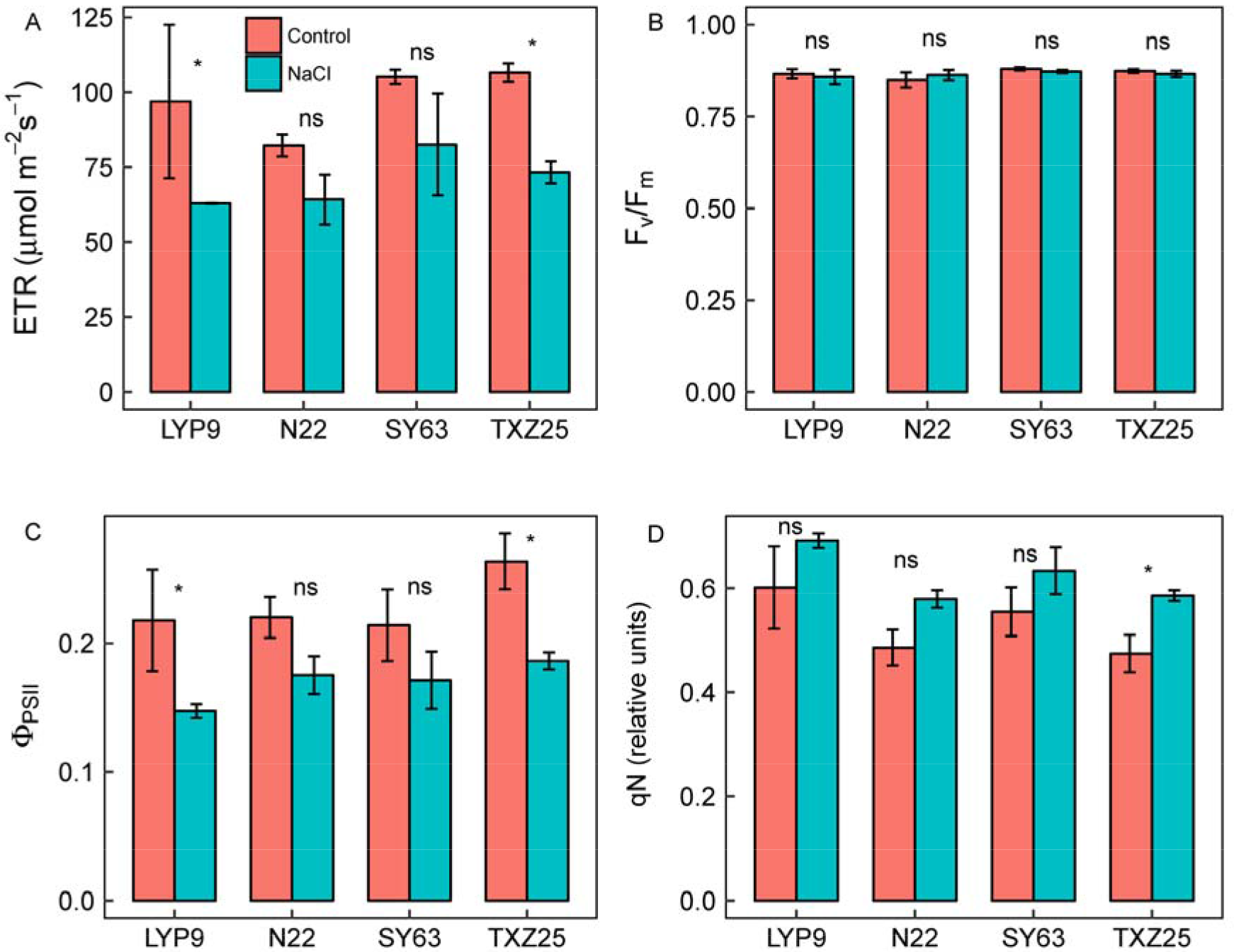
Effects of 7-days salt treat on (A) electron transport rate (ETR), (B) the maximum quantum efficiency (Fv/Fm), (C) actual quantum efficiency (Φ_PSII_), and (D) non-photochemical quenching coefficient (qN) of four rice genotypes. Bars represent the mean ± se of at least three replicates. ns, no significant; *, *P*<0.05; **, *P*<0.01; and ***, *P*<0.001.

**Fig. 5.**
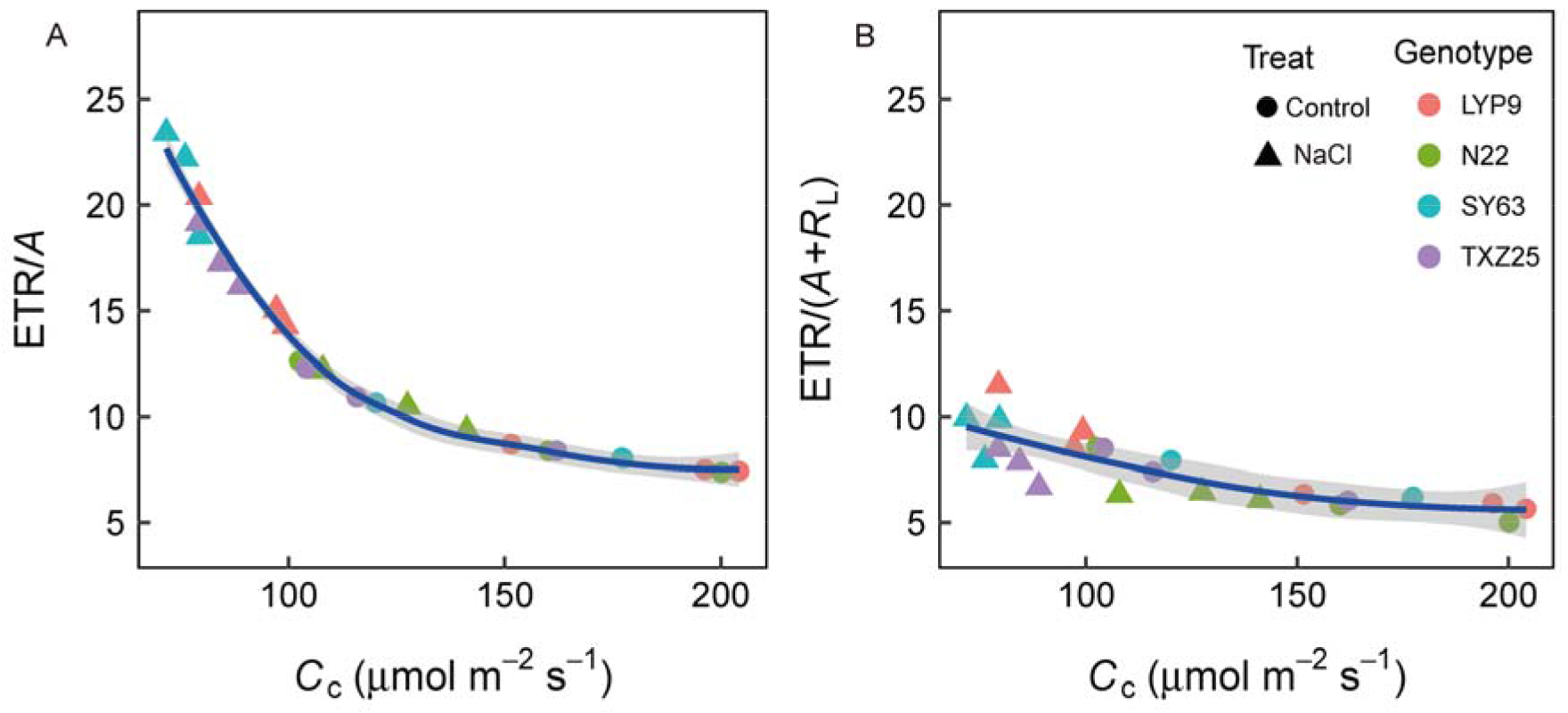
Ratio of electron transport rate (ETR) to (A) light-saturated photosynthetic rate (*A*) and to (B) gross CO_2_ assimilation accounting for photorespiration (*A*+*R*_L_) vs CO_2_ concentration in chloroplasts(*C*_c_)

### Limitation analysis

The impact of seven-days of salinity treatment on the relative stomatal (*l*_s_), mesophyll (*l*_m_) and biochemical (*l*_m_) limitations are shown in Fig. S4. Under normal condition (control) the *A* of the estimated rice genotypes was mainly limited by *l*_b_. However, in salt-stressed leaves, both *l*_s_ and *l*_m_ increased in all of the genotypes except the *l*_m_ in N22. In Fig. 6, the contributions of three relative limitations to decrease *A* are shown. In salt-stressed leaves, *L*_S_ (averaging 25%) and *L*_M_ (averaging 30%) increased dramatically in all four genotypes; however, the *L*_B_ was relatively small, except in N22.

**Fig. 6.**
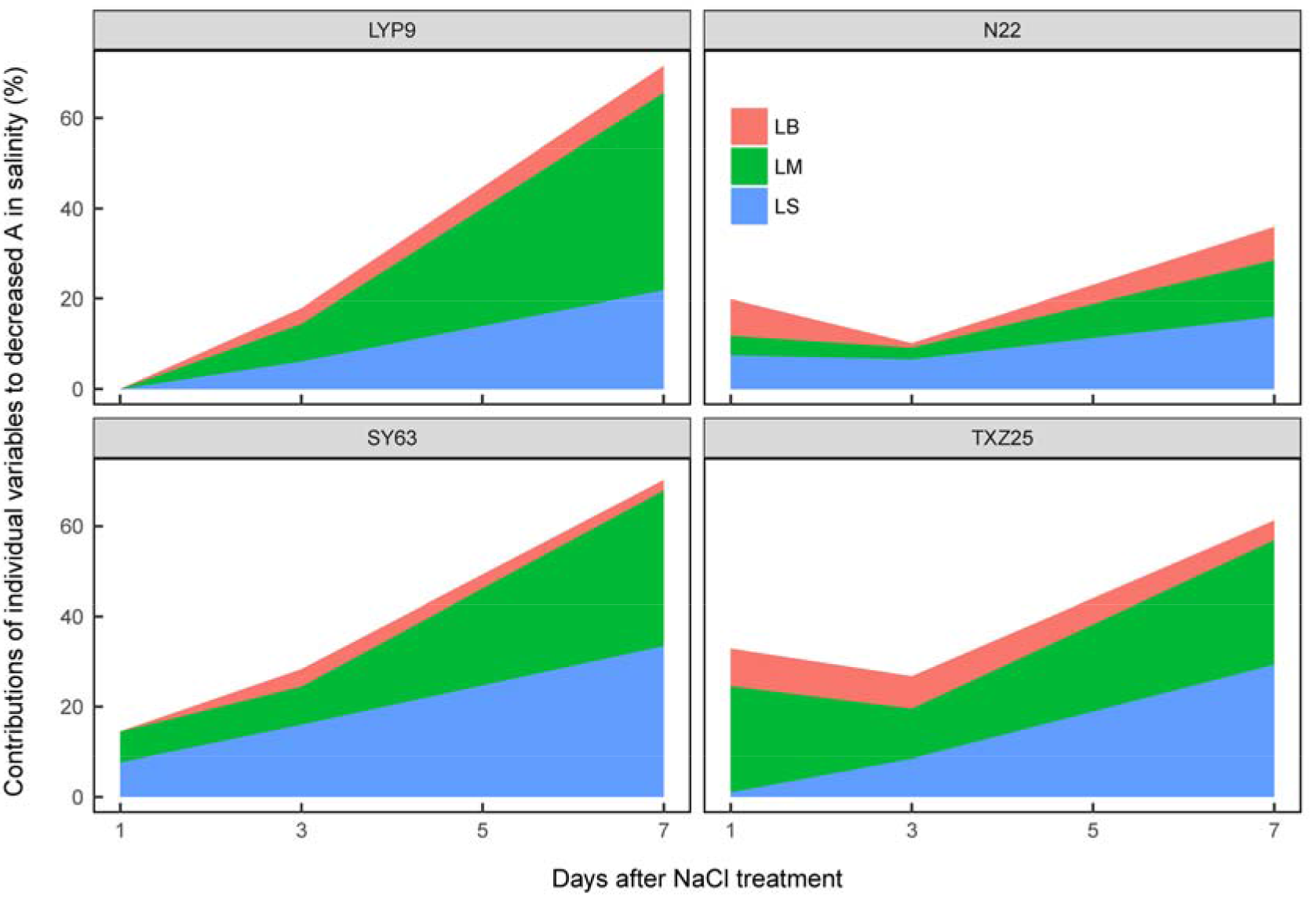
Contributions of stomatal conductance limitation (LS), mesophyll conductance limitation (LM) and biochemical limitation (LB) to decreases in light-saturated photosynthetic rate (*A*) of four rice genotypes.

Although both the leaf osmotic potential and Na^+^ content varied greatly among genotypes and NaCl treatments (Fig. S5, Table 2), the linear relationships between *l*_s_ and the leaf osmotic potential (*R*^2^=0.48; *P*=0.033) were found across genotypes and NaCl treatments, but not between *I*_s_ and the leaf Na^+^ content (Fig. 7). Moreover, the negative correlations between the Na^+^ content and *g*_s_ as well as transpiration rate (*E*) were found (Fig. S6). Unlike *l*_s_, lm linearly correlated with the leaf osmotic potential (*R*^2^=0.86; *P*<0.001) and leaf Na^+^ content (*R*^2^=0.52; *P*=0.026).

**Fig. 7.**
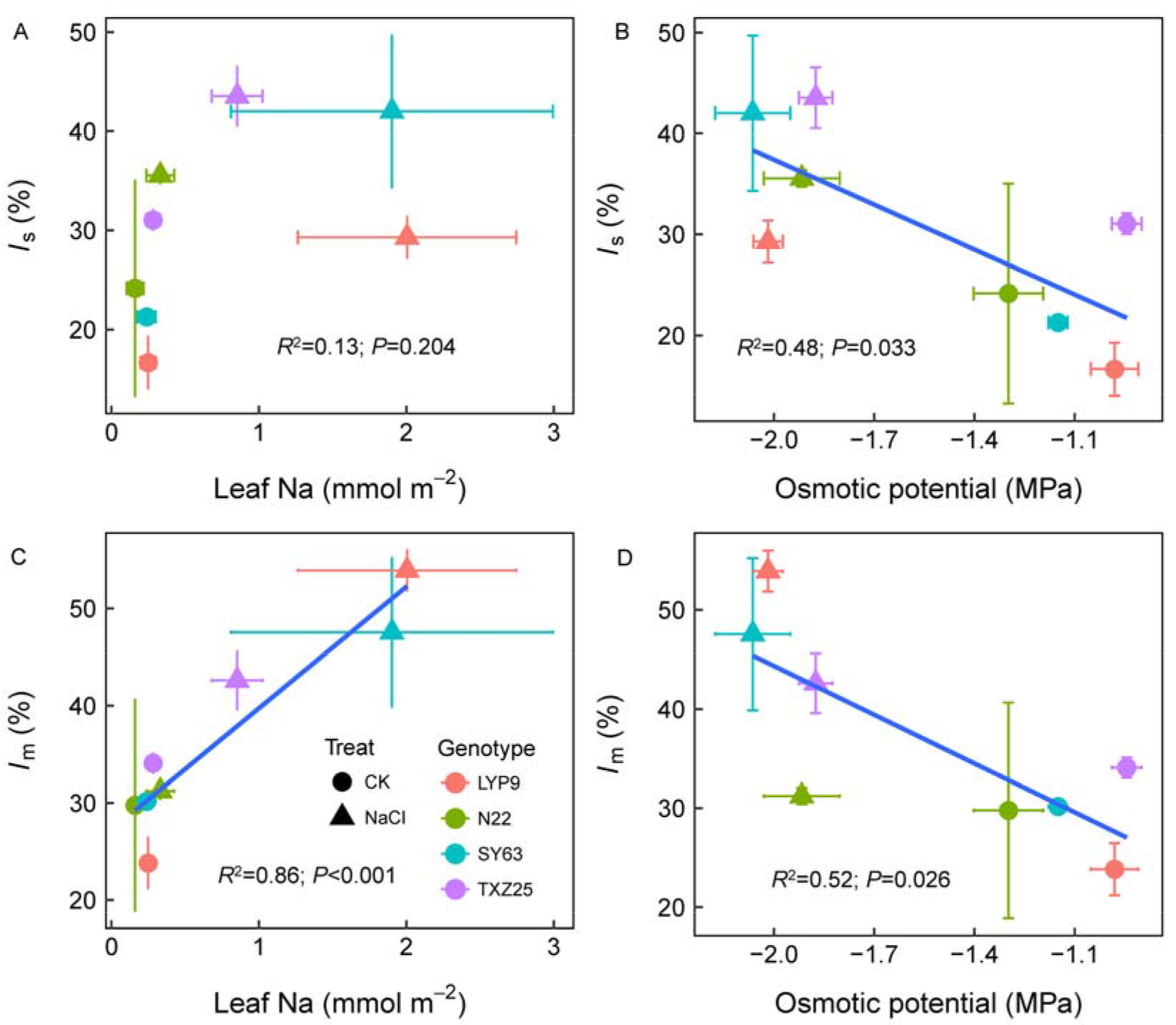
Effects of leaf Na content and osmotic potential on stomatal limitation (*l*_s_) (A, B) and mesophyll limitation in four rice genotypes. Points represent the mean ± se of at least three replicates.

## Discussion

### What determines the CO_2_ assimilation rate of salt-stressed leaves in rice?

In the present study, we showed that the leaf physiological and biochemical traits of rice were dramatically affected by soil salinity (Table 1, 2; Fig. 1-3). After seven days of NaCl treatment, the *A* decreased significantly in LYP9, SY63 and TXZ25 but not in N22. Generally, it is assumed that stomatal closure is the first response to salinity due to osmotic stress (Centritto et al. 2003, Chaves et al. 2011, Chen et al. 2015, Delfine et al. 1999, Delfine et al. 1998, Moradi and Ismail 2007). However, we observed that *g*_s_, *g*_m_ and ETR decreased dramatically in some of the genotypes one day after NaCl treatment, and multiple leaf parameters involving biochemical and physiological traits were affected in almost all of the genotypes after seven days of treatment (Fig. 1). To quantify the stomatal, mesophyll and biochemical limitations on *A* in salt-stressed rice leaves, the limitation analysis approach was used here. The results highlighted that CO_2_ diffusion conductance from the atmosphere to the sites of carboxylation (*g*_s_ and *g*_m_) played a key role in limiting *A* under salt stress (Fig. 6; Fig. S3), whereas biochemical factors played an important role in limiting *A* in rice under normal conditions (Fig. S3 C). In contrast to the previous studies of *Cucumis sativus* (Chen et al. 2015) and *Hordeum vuliare* (Perez-Lopez et al. 2012), mesophyll limitation (LM) contributed largely to reducing *A* in salt-stressed rice leaves. In fact, *g*_m_ was not affected by salinity in *H. vulgare*, and only a slight change was observed in *C. sativus*; however, *g*_m_ decreased more than 50% in all the estimated rice genotypes, except N22 in the current study (Fig. 2). Indeed, the decline in *A* that occurred in the salt stressed rice leaves was closely correlated with the low *g*s and *g*_m_ (Fig. S2). The contributions of biochemical limitation (BL) to reducing *A* in salt-stressed rice leaves were relatively small (Fig. 6), in disagreement with two studies on *H. vulgare* (Perez-Lopez et al. 2012) and *C. sativus* (Chen et al. 2015) under salt stress. This might be explained by a lower *C*_c_ in the current study than in these other studies. In the current study, the *C*_c_ in the salt-stressed leaves was typically lower than 100 μmol mol^−1^, except in N22 (Table 2); however, in the study of Perez-Lopez et al. (2012) the *C*_c_ was higher than 140 μmol mol^−1^. In fact, when the *C*_c_ is relatively higher (i.e. under normal conditions) biochemical factors were the predominat photosynthetic limiting factors in rice (Fig. S4).

Our results indicate that the influences of salt stress on protein and Rubisco contents varied greatly between genotypes (Table 1). In fact, the degradation of Rubisco-the most abundant protein-in the process of forming chloroplast protrusions (CPs) in salt-stressed rice has been observed in a previous study (Yamane et al. 2012). Moreover, the response of the Rubisco content to salt stress was observed to be dependent on salt treatment duration (Delfine et al. 1998). Therefore, the strong decrease in Rubisco content in TXZ25 and N22 might indicate a fast degradation in those genotypes. Although the Rubisco content decreased in N22 and TXZ25, the decline in *V*_cmax_ in salt-stressed leaves was relatively small, which supported the hypothesis that Rubisco also plays a role as a storage protein in C_3_ plant and is a major source of nitrogen for remobilization (Masclaux-Daubresse et al. 2010, Sage et al. 1987). More importantly, the apparent Rubisco activity (CE) was not affected by salt stress in rice, which also supports the idea that biochemical traits may not be the key factor causing decrease *A* in salt-stressed leaves (Table 2). Overall, the results indicate that the reduction of *A* in salt-stressed rice leaves was mainly related to the low *g*_s_ and *g*_m_ under salt stress.

### Salinity effects on CO_2_ diffusion

The *g*_s_ was determined by both stomatal anatomy (i.e., size and density) and opening status under given ambient air conditions (Xiong et al. 2017). While we did not investigate stomatal anatomical traits, it is unlikely that the stomatal size and density of the fully expand leaves can change fast enough to explain the decline in *g*_s_ by salinity over a very short time in the present study. Many previous studies (Centritto et al. 2003, Chaves et al. 2011, Chen et al. 2015, Delfine et al. 1999, Delfine et al. 1998, Moradi and Ismail 2007) have reported that osmotic stress caused by salinity can decrease the leaf osmotic/water potential, and then provoke stomatal close. Moreover, ionic stress due to the high leaf-Na^+^ content has been suggested as another factor provoking stomatal closure in *Aster tripolium* (Perera et al. 1994). The regressive analysis showed that stomatal limitation correlated with leaf osmotic potential but not with leaf Na^+^ content in rice (Fig. 7). The results suggest that salinity induced low *g*_s_ is mainly related to osmotic stress rather than ion stress in rice.

In general, *g*_m_ is related to leaf anatomical and biochemical traits under a given measurement condition (Evans et al. 2009, Tomas et al. 2013, Xiong et al. 2017). Previous studies have shown that long-term salinity significantly influences the leaf anatomical traits (Delfine et al. 1998, Wankhade et al. 2013); however, short-term salt-stress as in this study, causing leaf anatomical variation has rarely been estimated. Generally, the cell wall thickness (*T*_cw_) and the chloroplast surface area facing the intercellular airspace per unit leaf area (*S*_c_) are the two most important parameters related to g_m_ (Evans et al. 2009, Tomas et al. 2013, Xiong et al. 2017). The changes in *T*_cw_ are one of the potential reasons for the decline in *g*_m_ under salt stress. This is because osmatic stress usually introduced changes in the bulk elastic modulus, which relates to the alternation of biochemical composition and/or the thickness of the cell wall (Flexas and Diaz-Espejo 2014). The *S*_c_ is related to the mesophyll cell shape and the chloroplast shape as well as the light-dependent chloroplast arrangement (movement) inside the cells. Chloroplast movement is believed to alleviate photodamage to photosystems under stress conditions and rapid rearrangement of chloroplasts can profoundly impact *S*_c_ (Tholen et al. 2008, Xiong et al. 2015a). Moreover, previous studies have shown that mesophyll and chloroplast shape can be dramatically affected by short-term dehydration (Scoffoni et al. 2017, Scoffoni et al. 2016), which indicates that the low *S*_c_ caused by osmotic stress might be one of the reasons for the low *g*_m_ in salt-stressed leaves. Indeed, the linear correlation between *l*_m_ and leaf osmotic potential was observed in the present study (Fig. 7). The effects of biochemical traits on *g*_m_ in salinity have been suggested as being related to the functions of AQPs on membranes and carbonic anhydrase (CA) in cytosol and chloroplast stroma (Gao et al. 2010, Hu et al. 2012, Pongsomboon et al. 2009).

Interestingly, the relationship (*R*^2^=0.86; *P* < 0.001) between *l*_m_ and the leaf Na^+^ content across rice genotypes and NaCl treatments was closer than the relationship (*R*^2^=0.52; *P*=0.026) between *l*_m_ and the osmotic potential. Moreover, although the osmotic potential in the salt-stressed leaves of N22 decreased significantly, the leaf Na^+^ content did not increase (Fig. S5). Surprisingly, the *g*_m_ and *l*_m_ in the salt-stressed leaves of N22 did not change. These results suggest that the decreased *g*_m_ in the salt-stressed rice leaves would be more related to the accumulation of Na^+^ (ion effect). However, the mechanisms of ion impacts on *g*_m_ are unclear. One possibility is that AQPs regulate the Na^+^ absorption/distribution and CO_2_ diffusion across membranes under salinity. Indeed, the positive effects of AQPs on *g*_m_ have been demonstrated by a large number of previous studies (Flexas et al. 2006, Galmés et al. 2007, Hanba et al. 2004, Mori et al. 2014, Uehlein et al. 2012, Yang et al. 2000). Recently, Gao et al. (2010) reported that overexpressing an AQP gene from wheat caused the transgenic Arabidopsis to have lower Na^+^ levels than WT plants under salt stress. Otherwise, the Cl^−^ concentration which often correlates linearly with Na^+^ concentration in salt stressed leaves (Chen et al. 2015), might be another important ions that affects *g*_m_ (Tavakkoli et al. 2011). Further studies providing insights into the impacts of salinity on mesophyll anatomy, AQPs and CA, and thus *g*_m_, are necessary.

### Salinity effects on leaf photochemistry and photorespiration

When both *g*_s_ and *g*_m_ decreased significantly, the salinity could be expected to reduce *C*_c_ in the leaves. We observed a strong decrease in *C*_c_ in salt-stressed rice leaves except in N22. When *A* is limited by a low *C*_c_, it has been suggested that an imbalance occurs between PSII photochemical activity and the electron requirement for photosynthesis; consequently, photoinhibition results (James et al. 2006, Munns et al. 2016, Perez-Lopez et al. 2012, Zhang et al. 2010). While qN increased slightly in salt-stressed leaves, F_v_/F_m_ was unaffected (Fig. 4), which suggests that permanent photoinhibition was not the key factor in decreasing *A*, as already observed in previous studies. Although the reduction in ETR and Φ_PSII_ was observed in rice plant (Fig. 4), as reported in many species by previous studies (Moradi and Ismail 2007, Stepien and Johnson 2009), the decrease in *A* was far more serious than ETR and Φ_PSII_ (Fig. 1). Indeed, when *C*_c_ was lower than 100 μmol mol^−1^, the ETR/A increased very fast with decreasing *C*_c_ in rice (Fig. 5A). The results indicate that alternative sinks (i.e., photorespiration and Mehler reaction) for electrons replaced photosynthesis. However, ETR/(*A*+*R*_L_) was almost constant with changes *C*_c_ in rice (Fig. 5B), although a slight increase was found at a very low *C*_c_. Our results suggest that most of the thylakoid electron transport in rice leaves was used for the carboxylation plus oxygenation of Rubisco, and alternative sinks for electrons such as the Mehler reaction (photoinhibition), were very low under salinity conditions. Interestingly, similar conclusions were drawn by Flexas et al. (2002) in grapevines under field drought conditions. In fact, the high O_2_/CO_2_ ratio inside chloroplasts that were caused by low *C*_c_ led to higher photorespiration in salt-stressed leaves.

### Conclusions

The present study clearly validated that the *A* in rice leaves is predominately limited by low *C*_c_ rather than biochemical factors (i.e., Rubisco activity) under salt stress. Low osmotic potential introduced by salinity caused a strong increase in stomatal limitation and mesophyll limitation, and the accumulation of ions enhanced the mesophyll limitation. The results of the present study suggest that photoinhibition does not seriously increase in salt-stressed leaves. Furthermore, N22 exhibited higher salt tolerance than the other three genotypes, because N22 can maintain a lower tissues Na^+^ concentrations and higher osmotic potential than other genotypes under the same soil salt concentrations.

### Author contributions

D.X. planned and designed the research; X.W. and W.W. performed the experiments; D.X. and X.W. analyzed the data; X.W., D.X., J.H. and S.P. wrote the manuscript.

## Acknowledgements

The authors thank Miss Qiuqian Hu for technique assistance in chlorophyll fluorescence measurement. This work was supported by National Natural Science Foundation of China (No. 31671620), the Program of Introducing Talents of Discipline to Universities in China (no. B14032), the Program for Changjiang Scholars and Innovative Research Team in University of China (IRT1247), the National High Technology Research and Development Program of China (no. 2014AA10A605) and the Bill and Melinda Gates Foundation (OPP51587)

## Supporting Information

Additional Supporting Information may be found in the online version of this article:

**Fig. S1.** (A) Relationship between photochemical efficiency of photosystem II (Φ_PSII_) and Φ_CO2_and (B) The relationship between mesophyll diffusion conductance (*g*_m_) measured with Harley’s method and with Ethier’s method.

**Fig. S2.** Changes of light-saturated photosynthetic rate (*A*), electron transport rate (ETR), stomatal conductance (*g*_s_) and mesophyll conductance (*g*_m_) to NaCl treatment time.

**Fig. S3.** Correlations of light-saturated photosynthetic rate (*A*) and (A) stomatal conductance (*g*_s_) and (B) mesophyll conductance (*g*_m_) across four rice genotypes.

**Fig. S4.** Effects of salinity on photosynthetic limitations of four rice genotypes. *l*_s_, stomatal limitation; *l*_m_, mesophyll limitation; and *l*_b_, biochemical limitation. Bars represent the means±se of at least three replicates.

**Fig. S5.** Impacts of salinity on (A) leaf osmotic potential and (B) leaf Na content in rice.

**Fig. S6.** Impacts of leaf Na content on (A) stomatal conductance to CO_2_ (*g*_s_) and (B) transpiration rate (*E*) across four rice genotypes.

## References

Buckley TN and Diaz-Espejo A (2014) Partitioning changes in photosynthetic rate into contributions from different variables. Plant Cell Environ 38: 1200–1211

Centritto M, Loreto F and Chartzoulakis K (2003) The use of low CO_2_ to estimate diffusional and non-diffusional limitations of photosynthetic capacity of salt-stressed olive saplings. Plant Cell Environ 26: 585–594

Chaves MM, Miguel Costa J and Madeira Saibo NJ (2011) Recent advances in photosynthesis under drought and salinity. Adv Bot Res 57: 49–104

Chen T-W, Kahlen K and StÜTzel H (2015) Disentangling the contributions of osmotic and ionic effects of salinity on stomatal, mesophyll, biochemical and light limitations to photosynthesis. Plant Cell Environ 38: 1528–1542

Delfine S, Alvino A, Villani MC and Loreto F (1999) Restrictions to carbon dioxide conductance and photosynthesis in Spinach Leaves recovering from salt stress. Plant Physiol 119: 1101–1106

Delfine S, Alvino A, Zacchini M and Loreto F (1998) Consequences of salt stress on conductance to CO_2_ diffusion, Rubisco characteristics and atomy of spinach leaves. Funct Plant Biol 25: 395–402

Escalona JM, Flexas J and Medrano H (1999) Stomatal and non-stomatal limitations of photosynthesis under water stress in field-grown grapevines. Funct Plant Biol 26: 421–433

Ethier GJ and Livingston NJ (2004) On the need to incorporate sensitivity to CO_2_ transfer conductance into the Farquhar-von Caemmerer-Berry leaf photosynthesis model. Plant Cell Environ 27: 137–153

Evans JR, Kaldenhoff R, Genty B and Terashima I (2009) Resistances along the CO_2_ diffusion pathway inside leaves. J Exp Bot 60: 2235–2248

Farquhar G, von Caemmerer Sv and Berry J (1980) A biochemical model of photosynthetic CO_2_ assimilation in leaves of C3 species. Planta 149: 78–90

Flexas J, Barbour MM, Brendel O, Cabrera HM, Carriquí M, Díaz-Espejo A, Douthe C, Dreyer E, Ferrio JP, Gago J, Gallé A, Galmés J, Kodama N, Medrano H, Niinemets Ü, Peguero-Pina JJ, Pou A, Ribas-Carbó M, Tomás M, Tosens T and Warren CR (2012) Mesophyll diffusion conductance to CO_2_: An unappreciated central player in photosynthesis. Plant Science 193-194: 70–84

Flexas J, Baron M, Bota J, Ducruet JM, Galle A, Galmes J, Jimenez M, Pou A, Ribas-Carbo M, Sajnani C, Tomas M and Medrano H (2009) Photosynthesis limitations during water stress acclimation and recovery in the drought-adapted Vitis hybrid Richter-110 (V. berlandierixV. rupestris). J Exp Bot 60: 2361–2377

Flexas J, Bota J, Escalona JM, Sampol B and Medrano H (2002) Effects of drought on photosynthesis in grapevines under field conditions an evaluation of stomatal and mesophyll limitations. Funct Plant Biol 29: 461–471

Flexas J and Diaz-Espejo A (2014) Interspecific differences in temperature response of mesophyll conductance: food for thought on its origin and regulation. Plant Cell Environ 38: 625–628

Flexas J, Díaz-Espejo A, Berry J, Cifre J, Galmés J, Kaldenhoff R, Medrano H and Ribas-Carbó M (2007) Analysis of leakage in IRGA's leaf chambers of open gas exchange systems: quantification and its effects in photosynthesis parameterization. J Exp Bot 58: 1533–154316

Flexas J, Diaz-Espejo A, Conesa MA, Coopman RE, Douthe C, Gago J, Galle A, Galmes J, Medrano H, Ribas-Carbo M, Tomas M and Niinemets U (2016) Mesophyll conductance to CO_2_ and Rubisco as targets for improving intrinsic water use efficiency in C_3_ plants. Plant Cell Environ 39: 965–982

Flexas J, Ribas-Carbo M, Hanson DT, Bota J, Otto B, Cifre J, McDowell N, Medrano H and Kaldenhoff R (2006) Tobacco aquaporin NtAQP1 is involved in mesophyll conductance to CO_2_ in vivo. Plant J 48: 427–439

Galle A, Florez-Sarasa I, Aououad HE and Flexas J (2011) The Mediterranean evergreen Quercus ilex and the semi-deciduous Cistus albidus differ in their leaf gas exchange regulation and acclimation to repeated drought and re-watering cycles. J Exp Bot 62: 5207–5216

Galle A, Florez-Sarasa I, Tomas M, Pou A, Medrano H, Ribas-Carbo M and Flexas J (2009) The role of mesophyll conductance during water stress and recovery in tobacco (Nicotiana sylvestris): acclimation or limitation? J Exp Bot 60: 2379–2390

Galmés J, Pou A, Alsina MM, Tomas M, Medrano H and Flexas J (2007) Aquaporin expression in response to different water stress intensities and recovery in Richter-110 (Vitis sp.): relationship with ecophysiological status. Planta 226: 671–681

Gao Z, He X, Zhao B, Zhou C, Liang Y, Ge R, Shen Y and Huang Z (2010) Overexpressing a putative aquaporin gene from wheat, TaNIP, enhances salt tolerance in transgenic Arabidopsis. Plant Cell Physiol 51: 767–775

Grassi G and Magnani F (2005) Stomatal, mesophyll conductance and biochemical limitations to photosynthesis as affected by drought and leaf ontogeny in ash and oak trees. Plant Cell Environ 28: 834–849

Hanba YT, Shibasaka M, Hayashi Y, Hayakawa T, Kasamo K, Terashima I and Katsuhara M (2004) Overexpression of the barley aquaporin HvPIP2;1 increases internal CO_2_ conductance and CO_2_ assimilation in the leaves of transgenic rice plants. Plant Cell Physiol 45: 521–529

Harley PC, Loreto F, Di Marco G and Sharkey TD (1992) Theoretical considerations when estimating the mesophyll conductance to CO_2_ flux by analysis of the response of photosynthesis to CO_2_. Plant Physiol 98: 1429–1436

Hermida-Carrera C, Kapralov MV and Galmes J (2016) Rubisco catalytic properties and temperature response in crops. Plant Physiol 171: 2549–2561

Hu W, Yuan Q, Wang Y, Cai R, Deng X, Wang J, Zhou S, Chen M, Chen L, Huang C, Ma Z, Yang G and He G (2012) Overexpression of a wheat aquaporin gene, TaAQP8, enhances salt stress tolerance in transgenic tobacco. Plant Cell Physiol 53: 2127–2141

James RA, Munns R, Von Caemmerer S, Trejo C, Miller C and Condon T (2006) Photosynthetic capacity is related to the cellular and subcellular partitioning of Na^+^, K^+^ and Cl^−^ in salt-affected barley and durum wheat. Plant Cell Environ 29: 2185–2197

James RA, Rivelli AR, Munns R and Caemmerer Sv (2002) Factors affecting CO_2_ assimilation, leaf injury and growth in salt-stressed durum wheat. Funct Plant Biol 29: 1393–1403

Khan HA, Siddique KHM, Munir R and Colmer TD (2015) Salt sensitivity in chickpea: Growth, photosynthesis, seed yield components and tissue ion regulation in contrasting genotypes. J Plant Physiol 182: 1–12

Koyro H-W, Hussain T, Huchzermeyer B and Khan MA (2013) Photosynthetic and growth responses of a perennial halophytic grass Panicum turgidum to increasing NaCl concentrations. Environ Exp Bot 91: 22–29

Masclaux-Daubresse C, Daniel-Vedele F, Dechorgnat J, Chardon F, Gaufichon L and Suzuki A (2010) Nitrogen uptake, assimilation and remobilization in plants: challenges for sustainable and productive agriculture. Ann Bot 105: 1141–1157

Moradi F and Ismail AM (2007) Responses of photosynthesis, chlorophyll fluorescence and ROS-scavenging systems to salt stress during seedling and reproductive stages in rice. Ann Bot 99: 1161–1173

Mori IC, Rhee J, Shibasaka M, Sasano S, Kaneko T, Horie T and Katsuhara M (2014) CO_2_ transport by PIP2 aquaporins of barley. Plant Cell Physiol 55: 251–257

Munns R, James RA, Gilliham M, Flowers TJ and Colmer TD (2016) Tissue tolerance: an essential but elusive trait for salt-tolerant crops. Funct Plant Biol 43: 1103–1113

Murchie EH and Lawson T (2013) Chlorophyll fluorescence analysis: a guide to good practice and understanding some new applications. J Exp Bot 64: 3983–3998

Negrão S, Courtois B, Ahmadi N, Abreu I, Saibo N and Oliveira MM (2011) Recent updates on salinity stress in rice: From physiological to molecular responses. Crit Rev Plant Sci 30: 329–377

Perera LKRR, Mansfield TA and Malloch AJC (1994) Stomatal responses to sodium ions in Aster tripolium: a new hypothesis to explain salinity regulation in above-ground tissues. Plant Cell Environ 17: 335–340

Perez-Lopez U, Robredo A, Lacuesta M, Mena-Petite A and Munoz-Rueda A (2012) Elevated CO_2_ reduces stomatal and metabolic limitations on photosynthesis caused by salinity in Hordeum vulgare. Photosynth Res 111: 269–283

Pongsomboon S, Udomlertpreecha S, Amparyup P, Wuthisuthimethavee S and Tassanakajon A (2009) Gene expression and activity of carbonic anhydrase in salinity stressed Penaeus monodon. Comp Biochem Physiol A Mol Integr Physiol 152: 225–233

Sade N, Gallé A, Flexas J, Lerner S, Peleg G, Yaaran A and Moshelion M (2014) Differential tissue-specific expression of NtAQP1 in Arabidopsis thaliana reveals a role for this protein in stomatal and mesophyll conductance of CO_2_ under standard and salt-stress conditions. Planta 239: 357–366

Sage RF, Pearcy RW and Seemann JR (1987) The Nitrogen use efficiency of C_3_ and C_4_ plants: III. Leaf nitrogen effects on the activity of carboxylating enzymes in Chenopodium album (L.) and Amaranthus retroflexus (L.). Plant Physiol 85: 355–359

Scoffoni C, Albuquerque C, Brodersen C, Townes SV, John GP, Bartlett MK, Buckley TN, McElrone AJ and Sack L (2017) Outside-xylem vulnerability, not xylem embolism, controls leaf hydraulic decline during dehydration. Plant Physiol 173: 1197–1210

Scoffoni C, Albuquerque C, Brodersen CR, Townes SV, John GP, Cochard H, Buckley TN, McElrone AJ and Sack L (2016) Leaf vein xylem conduit diameter influences susceptibility to embolism and hydraulic decline. New Phytol 213: 1076–1092

Stepien P and Johnson GN (2009) Contrasting responses of photosynthesis to salt stress in the glycophyte Arabidopsis and the halophyte thellungiella: role of the plastid terminal oxidase as an alternative electron sink. Plant Physiol 149: 1154–1165

Tavakkoli E, Fatehi F, Coventry S, Rengasamy P and McDonald GK (2011) Additive effects of Na+ and Cl- ions on barley growth under salinity stress. J Exp Bot 62: 2189–2203

Tholen D, Boom C, Noguchi K, Ueda S, Katase T and Terashima I (2008) The chloroplast avoidance response decreases internal conductance to CO_2_ diffusion in Arabidopsis thaliana leaves. Plant Cell Environ 31: 1688–1700

Tomas M, Flexas J, Copolovic iL, Galmes J, Hallik L, Medrano H, Ribas-Carbo M, Tosens T, Vislap V and Niinemets U (2013) Importance of leaf anatomy in determining mesophyll diffusion conductance to CO_2_ across species: quantitative limitations and scaling up by models. J Exp Bot 64: 2269–2281

Tosens T, Nishida K, Gago J, Coopman RE, Cabrera HM, Carriqui M, Laanisto L, Morales L, Nadal M, Rojas R, Talts E, Tomas M, Hanba Y, Niinemets U and Flexas J (2015) The photosynthetic capacity in 35 ferns and fern allies: mesophyll CO_2_ diffusion as a key trait. New Phytol 209: 1576–1590

Uehlein N, Sperling H, Heckwolf M and Kaldenhoff R (2012) The Arabidopsis aquaporin PIP1;2 rules cellular CO_2_ uptake. Plant Cell Environ 35: 1077–1083

Valentini R, Epron D, De Angelis P, Matteucci G and Dreyer E (1995) In situ estimation of net CO_2_ assimilation, photosynthetic electron flow and photorespiration in Turkey oak (Q. cerris L.) leaves: diurnal cycles under different levels of water supply. Plant Cell Environ 18: 631–640

Wankhade SD, Cornejo M-J, Mateu-Andrés I and Sanz A (2013) Morpho-physiological variations in response to NaCl stress during vegetative and reproductive development of rice. Acta Physiol Plant 35: 323–333

Xiong D, Chen J, Yu T, Gao W, Ling X, Li Y, Peng S and Huang J (2015a) SPAD-based leaf nitrogen estimation is impacted by environmental factors and crop leaf characteristics. Sci Rep 5: 13389

Xiong D, Flexas J, Yu T, Peng S and Huang J (2017) Leaf anatomy mediates coordination of leaf hydraulic conductance and mesophyll conductance to CO_2_ in Oryza. New Phytol 213: 572–583

Xiong D, Liu X, Liu L, Douthe C, Li Y, Peng S and Huang J (2015b) Rapid responses of mesophyll conductance to changes of CO_2_ concentration, temperature and irradiance are affected by N supplements in rice. Plant Cell Environ 38: 2541–2550

Xiong D, Yu T, Liu X, Li Y, Peng S and Huang J (2015c) Heterogeneity of photosynthesis within leaves is associated with alteration of leaf structural features and leaf N content per leaf area in rice. Funct Plant Biol 42: 687–696

Yamane K, Mitsuya S, Taniguchi M and Miyake H (2012) Salt-induced chloroplast protrusion is the process of exclusion of ribulose-1,5-bisphosphate carboxylase/oxygenase from chloroplasts into cytoplasm in leaves of rice. Plant Cell Environ 35: 1663–1671

Yang B, Fukuda N, van Hoek A, Matthay MA, Ma T and Verkman A (2000) Carbon dioxide permeability of aquaporin-1 measured in erythrocytes and lung of aquaporin-1 null mice and in reconstituted proteoliposomes. J Biol Chem 275: 2686–2692

Zhang T, Gong H, Wen X and Lu C (2010) Salt stress induces a decrease in excitation energy transfer from phycobilisomes to photosystem II but an increase to photosystem I in the cyanobacterium Spirulina platensis. J Plant Physiol 167: 951–958

